# Individual differences in members of Actinobacteria, Bacteroidetes, and Firmicutes is associated with resistance or vulnerability to addiction-like behaviors in heterogeneous stock rats

**DOI:** 10.1101/2021.07.23.453592

**Authors:** S. Simpson, G. de Guglielmo, M. Brennan, L. Maturin, G. Peters, H. Jia, E. Wellmeyer, S. Andrews, L. Solberg Woods, A. A. Palmer, O. George

**Author notes:** Corresponding Author <u>Dr. Olivier George,</u>, Department of Psychiatry, University of California San Diego La Jolla, CA 92093, USA.

## Abstract

An emerging element in psychiatry is the gut-brain-axis, the bi-directional communication pathways between the gut microbiome and the brain. A prominent hypothesis, mostly based on preclinical studies, is that individual differences in the gut microbiome composition and drug-induced dysbiosis may be associated with vulnerability to psychiatric disorders including substance use disorder. However, most studies used small sample size, ignored individual differences, or used animal models with limited relevance to addiction. Here, we test the hypothesis that pre-existing microbiome composition and drug-induced changes in microbiome composition can predict addiction-like behaviors using an advanced animal model of extended access to cocaine self-administration in a large cohort of heterogenous stock (HS) rats. Adult male and female HS rats were allowed to self-administer cocaine under short (2h/day) and long access (6h/day) for ~7 weeks under various schedule of reinforcement to identify individuals that are resistant or vulnerable to addiction-like behaviors and fecal samples were collected before the first session and after the last session to assess differences in the microbiome composition. Linear discriminant analysis (LDA) identified sex-dependent and sex-independent differences at the phylum, order, and species level that are differentially abundant in resistant vs. vulnerable individuals, including high level of actinobacteria both before the first exposure to cocaine and after 7 weeks of cocaine self-administration in resistant animals. Predictions of functional gene content using PICRUSt revealed differential regulation of short-chain fatty acid processing in the vulnerable group after self-administration. These results identify microbiome constituents as well as metabolic pathways that are associated with resistance or vulnerability to addiction-like behaviors in rats. Identification of microbes and tangential metabolic pathways involved in cocaine resilience/vulnerability may represent an innovative strategy for the development of novel biomarkers and medication for the treatment of cocaine use disorder.

## Introduction

Despite the fall in popularity of cocaine after the 1980’s, cocaine use disorder (CUD) continues to be a major public health problem in the United States. Over 2 million people are estimated to have CUD in the US and during the current COVID crisis, cocaine use and cocaine related overdose deaths have multiplied (Wainwright et al., 2020; SAMHSA, 2017). Currently, there are no approved medications for the treatment of cocaine use disorder and the factors that contribute to CUD are not well understood (Cano Fischer et al., 2015; Simpson et al., 1999. Genetics, socioeconomic status, and other outside factors are known to play a role in addiction liability; novel approaches to investigate contributing factors are necessary to better understand and treat CUD.

In the past decade, the gut-brain axis has garnered attention for the potential involvement of the microbiome in substance use disorders (Kiraly et al., 2016b; Meckel and Kiraly, 2019; Dinan and Cryan, 2017; Simpson et al., 2020a), comorbid psychiatric conditions (Kim and Shin, 2018; Cryan and Dinan, 2012; Diaz Heijtz et al., 2011), and other disease states (Tilg et al., 2016). Microbes communicate through the gut-brain axis through three major pathways 1) chemical signaling (metabolites), 2) physical connections (vagus nerve), and 3) multi-functional pathways (HPA Axis/immune system) (Cryan and Dinan, 2012; Cryan et al., 2019). Lifestyle and genetics contribute to differences observed in microbial communities in both animals and humans, which may have downstream consequences to mental health, addiction liability, and disease states.

Early work in animals has demonstrated that depletion of the microbiome using antibiotics alters cocaine conditioned place preference (CPP) to sub-reward threshold levels that is ameliorated by application of a cocktail of microbial metabolites known as short-chain fatty acids (SCFA). (Kiraly et al., 2016b). SCFA’s are a critical energy sources (LeBlanc et al., 2017) and anti-inflammatory molecules (Vinolo et al., 2011). SCFA’s signal in a variety of ways including through g-protein coupled receptors (GPCRs) (Kimura et al., 2011) and act as histone-deacetylase inhibitors (Fellows et al., 2018). Similar observations have been made in other stimulants such as methamphetamine where depletion of the microbiome by antibiotic perturbation altered CPP responses (Ning et al., 2017). In other drugs of abuse, depletion of the microbiome in opioids did not alter behavior, but alter the neuronal ensembles in states of intoxication and withdrawal (Simpson et al., 2020b). The severity of drug intake has been correlated with distinct microbiome profiles in alcoholics (Mutlu et al., 2012; Leclercq et al., 2017). Alcohol has also been demonstrated to differentially impact key microbiota in a sex specific manner known for immunoregulatory roles such as *Turcibacter* and *Clostridium* (Caslin et al., 2019). Psychiatric disorders that are often co-morbid (and contribute to) drug use have also been associated with alterations of the gut microbiome, such as depression (Limbana et al., 2020), anxiety (Yang et al., 2019), and related immunological states (Peirce and Alvina, 2019). These lines of evidence support that the microbiome as a rich target for the discovery of potential treatments either via direct microbial administration or downstream metabolites.

The present study tested the hypothesis that pre-drug microbiome states are stratified based on behavioral outcomes and that protective microbes could be identified in a model of genetically diverse animals that are translationally relevant and representative of the genetic diversity observed in human populations. Both preclinical and clinical investigation of the microbiome report differences in the microbiome related to sex and genetics (Kim et al., 2020; Goodrich et al., 2014), and fluctuations to the microbiome occur as we age (Jasarevic et al., 2016) and with hormone status (Valeri and Endres, 2021). To address these questions, we analyzed the microbiome differences between HS rats by addiction index (Kallupi et al., 2020), a multifaceted measure of cocaine use and by sex to identify microbial constituents that may contribute to resistance/vulnerability to escalation of cocaine use.

## Materials and Methods

### Animals

Adult female (n=27) and male (n=31) Heterogenous Stock (HS) rats were provided by Dr. Leah Solberg Woods (Wake Forest University School of Medicine). Rats were housed two per cage on a reverse 12h/12h light/dark cycle (lights off at 8:00 am) in a temperature (20–22°C) and humidity (45–55%) controlled animal facility with ad libitum access to water and food pellets (PJ Noyes Company). All procedures were conducted in adherence to the National Institutes of Health Guide for the Care and Use of Laboratory Animals and were approved by the Institutional Animal Care and Use Committee of The University of California San Diego.

### Drugs

Cocaine HCl (National Institute on Drug Abuse, Bethesda, MD) was dissolved in 0.9% saline (Hospira, Lake Forest, IL) at a dose of 0.5 mg/kg per 100ul infusion. Animals self-administered the drug intravenously via an external port connected to a jugular vein catheter.

### Intravenous catheterization

Rats were anesthetized with 1-5% isoflurane. Intravenous catheters were implanted in the right jugular vein. The catheter was constructed from an 18 cm length of Micro-Renathane tubing (0.023-inch inner diameter, 0.037-inch outer diameter; Braintree Scientific, Braintree, MA, USA) attached to a 90° guide cannula (Plastics One, Roanoke, VA, USA), which is embedded in dental acrylic, and anchored under the skin with a 2 cm square of mesh. The jugular vein was punctured with a 22-gauge needle, and then the tubing was inserted and secured inside the vein with suture thread. The catheter port is extended through an incision on the back and sealed with a plastic cap and metal cover. Catheters were flushed daily with heparinized saline (10 U/ml of heparin sodium; American Pharmaceutical Partners, Schaumburg, IL, USA) in 0.9% bacteriostatic sodium chloride (Hospira, Lake Forest, IL, USA) that contained 52.4 mg/0.2 ml of the antibiotic Cefazolin (Hospira, Lake Forest, IL, USA). Catheter patency is tested at the start and end of the experiment using a short-acting barbiturate.

### Behavioral Testing

Behavioral testing was performed during the animals’ dark cycle from age 7 weeks – 15 weeks. According to the Cocaine Biobank protocol (Carrette et al., 2021), self-administration was performed in Med Associates operant chambers (28×24×19.5cm) equipped with two retractable levers. Each session was started when the levers were extended into the chamber. Cocaine was delivered through an infusion pump that was activated by responses on the active lever (right), resulting in the delivery of cocaine (0.5 mg/kg per 100ul). Rewards were paired with a cute light upon activation of the active lever followed by a 20 second timeout period. Responses on the inactive lever (left) were recorded but did not have scheduled consequences. The rats were first trained to self-administer cocaine under short-access (ShA) fixed-ratio 1 (FR1) schedule of reinforcement in daily 2-h sessions for 10 days. After the ShA epoc was completed, they were then able to self-administer for long access sessions (LgA) for 6-h/ day for 14 days with weekend breaks (Ahmed and Koob, 1998).

### Progressive Ratio (PR)

At the end of each epoc (ShA and LgA phases), a progressive ratio (PR) schedule of reinforcement was used to measure of the reinforcing value of a reward (Stafford et al., 1998; Hodos, 1961). The PR response requirements necessary to receive a single drug dose increased according to this equation: [5e ^(injection numbers x 0.2)^] – 5, resulting in the following progression of response requirements: 1, 2, 4, 6, 9, 12, 15, 20, 25, 32, 40, 50, 62, 77, 95, 118, 145, 178, 219, 268, etc. The breakpoint was defined as the last ratio attained by the rat prior to a 60 min period during which a ratio was not completed, after which the program terminates the test. Progressive ratio testing was repeated two times after the LgA phase.

### Compulsive-like responding contingent footshock

After LgA sessions and PR testing, rats were placed in the self-administration chamber for 1 h and tested for compulsive-like behavior. During this protocol, 30% of the reinforced responses were paired with a contingent footshock (0.3 mA, 0.5 s).

### Feces collection

At times marked in Figure 1a (BSL/LGA), feces were collected for microbiome analysis. The anal canal was gently massaged to collect 2 fecal pellets which were immediately put into a sterile tube, and snap-frozen on dry ice, then stored at −80°C until processing. Feces collected at the LGA timepoint were taken 16-18h following the last session during the withdrawal phase.

**Figure 1:**
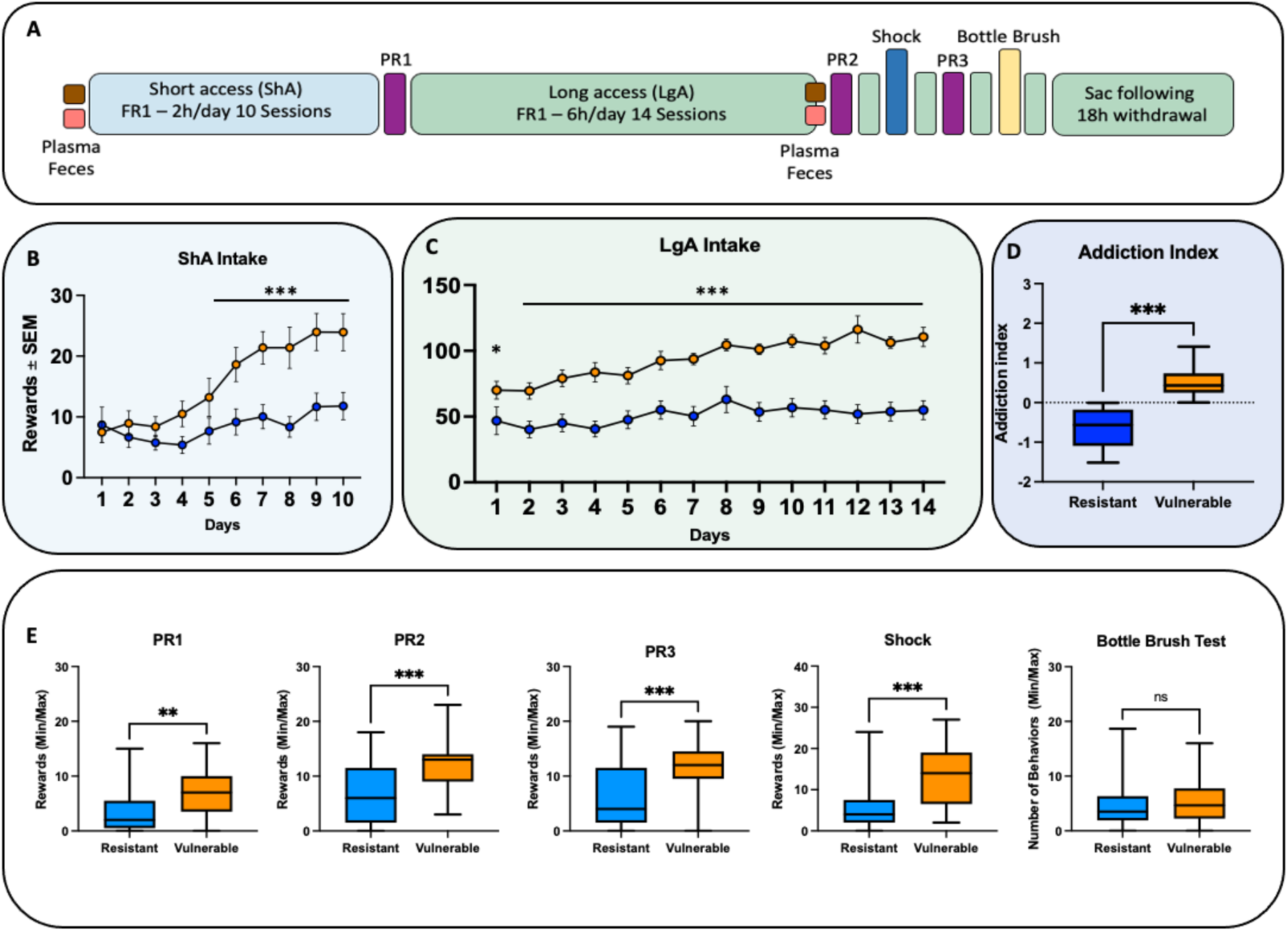
Experimental design and behavioral outcomes. **A)** Diagram of the experimental design for the timepoints of all behavioral procedures. Longitudinal samples (feces and plasma) were harvested before and after drug exposure (pink and brown boxes). Based on the addiction index **(D)**, animals were split at the median and assessed for cocaine rewards each day of short and long access. (**B)** During short access (ShA), a two-way ANOVA revealed a main effect (time x group) interaction (p < 0.001, F_(9,252)_ = 5.74), a Bonferroni post-hoc revealed increased rewards in the Vulnerable group compared to the Resistant group at days 6-10 (p < 0.001). **(C)** During long access (LgA), a two-way ANOVA revealed a main effect (time x group) interaction (p < 0.002, F_(13,364)_ = 2.539), a Bonferroni post-hoc revealed the Vulnerable group had more rewards at day 1 (p=0.012), and days 2-14 (p < 0.001) **(E)**. Resistant animals exhibited a lower number of rewards for all progressive ratio tests compared to vulnerable animals - PR1 (p = 0.008, t=2.757), PR2 (p < 0.001, t=4.351), PR3 was (p < 0.001, t=3.878) indicating reduced motivation for cocaine reward. Resistant animals were less likely to respond during the shock session demonstrating a lower level of compulsivity compared to vulnerable animals (p<0.001), t=4.285). There was no difference in irritability measures between the resistant and vulnerable groups when comparing the bottle brush irritability test (p=0.625, t=0.4907).

### 16S rRNA sequencing

At the time of DNA extraction, feces were thawed and extracted with the Qiagen DNeasy powersoil kit. The samples are then amplified by PCR in triplicate and then pooled. The 16S V4 gene was amplified using universal primers (525F806R). Each sample was normalized to 240 ng per sample and purified. After purification, the A260/A280 ratio of the final pool was recorded to ensure purity, with a tolerance range of 1.8–2.0. The barcoded amplicons from all samples were normalized, pooled to construct the sequencing library, and sequenced using an Illumina MiSeq Sequencer. The sequencing primers were the following: forward, TATGGTAATTGTGTGYCAGMGCCGCGGTAA and reverse, AGTCAGCCAGCCGGACTACNVGGGTWT CTAAT; and index sequence, AATGATACGGCGACCA CCGAGATCTACACGCT.

### 16s sequence analysis

16S rRNA gene sequence analysis was performed as previously described (Simpson et al., 2020b) After sequencing, the raw files were prepared, filtered for quality, and demultiplexed. Operational taxonomic units (OTUs) were selected using open reference OUT picking based on 97% sequence similarity to the Greengenes database (McDonald et al., 2012). Taxonomy assignment and rarefaction were performed using QIIME/Qiita with 9,000 reads per sample (Caporaso et al., 2012).

### PICRUSt

PICRUSt (v. 1.1.4) (Langille et al., 2013) creates functional predictions based on 16s sequencing profiles. Functional genes and pathways were identified using KEGG (Kanehisa and Goto, 2000). https://github.com/picrust/picrust.

### Statistics

Statistics on behavioral outcomes, including median splits and z-score calculations were conducted with PRISM 9 GraphPad (San Diego, California, USA). Statistical tests were performed as described. Post hoc tests were Bonferroni corrected. Linear discriminant analysis Effect Size (LEfSe) was accomplished using (http://huttenhower.sph.harvard.edu/galaxy/) (Goecks et al., 2010) with stringent cutoffs to filter only results (p<0.05).

## Results

### Resistant and vulnerable animals differ by self-administration and progressive ratio

Rats self-administered cocaine under the experimental timeline in **Figure 1a**. At the end of the experiment animals were split by an addiction index that created by an average of the z-scores of the escalation, progressive ratio, and shock responses. Subjects were split into “Resistant” and “Vulnerable” groups, using a median split of the addiction index. Intake at short access, long access, and measures for each behavioral task were compared. During the 2-hour short access self-administration sessions (ShA) **(Figure 1B)** a two-way ANOVA revealed a main effect (time x group) interaction (p < 0.001, F_(9,252)_ = 5.74), a Bonferroni post-hoc revealed increased rewards in the Vulnerable group compared to the Resistant group at days 6-10 (p < 0.001). During the 6-hours long access self-administration sessions (LgA) **(Figure 1C)** a two-way ANOVA revealed a main effect (time x group) interaction (p < 0.002, F_(13,364)_ = 2.539), a Bonferroni post-hoc revealed the Vulnerable group had more rewards at day 1 (p=0.012), and days 2-14 (p < 0.001) when compared to the Resistant group. The addiction index, plotted in **Figure 1D** was assessed by a two tailed students t-test (p <0.001), the mean of the resistant animals was −0.6124, and the mean of the vulnerable animals was 0.5173. Each behavioral test listed in **Figure 1A** was compared between resistant and vulnerable animals for PR1, PR2, PR3, Shock, and the Bottle Brush Irritability test. Resistant animals exhibited a lower number of rewards for all progressive ratio tests compared to vulnerable animals - PR1 (p = 0.008, t=2.757), PR2 (p < 0.001, t=4.351), PR3 was (p < 0.001, t=3.878) indicating reduced motivation for cocaine reward. Resistant animals were less likely to respond during the shock session demonstrating a lower level of compulsivity compared to vulnerable animals (p<0.001), t=4.285). There was no difference in irritability measures between the resistant and vulnerable groups when comparing the bottle brush irritability test (p=0.625, t=0.4907).

### Resistant and Vulnerable Populations Exhibit Differences in Microbiome Composition at Baseline

To assess the basal microbiome composition of the Resistant and Vulnerable animal groups, we analyzed microbiome differences at the phylum level **(Figure 2A)**. Proteobacteria (p=0.239), Firmicutes (p=0.307), Bacteroidetes (p>0.999), Verrucomicrobia (p=0.0778), Cyanobacteria (p=0.0228), Tenericutes (p=0.610), and Spiroachetes (p=0.245) were all not significantly different between the groups; however, Actinobacteria, the third largest phylum had a higher percent abundance in the Resistant group when compared to the Vulnerable group (p =0.025, Mann-Whitney U = 277). Stacked bar charts shows the representative individuals from each group **(Figure 2B)**, and donut plots show the average relative abundance of each group. The LEfSe taxonomic cladogram (**Figure 2C**) demonstrated increased abundance in the classes Actinobacteria and Bifidobacterium in the Resistant group, increased abundance in the order Bacteriodales in the Resistant group and Turcibacter in the Vulnerable group, increased abundance in the family, *Peptostreptococcacea, Bifidobacteriaceae* in the Resistant group, and *Staphylococcacacae*, *Turcibacteraceae, Odoribacteraceae* in the Vulnerable group, increased abundance in the genus *Bifidobacterales, Butyrucimonas, Allobaculum, and Sphingomonas* in the Resistant group, and increased abundance of *Turcibacterales, Sarcina, Dorea in the Vulnerable group*, as well increased abundance at the species level of *Manihotvorans, Areus* in the Resistant group, and *Eutactus*, in the Vulnerable group. The differential abundance is demonstrated in the LDA Score bar plots to the right of the cladogram.

**Figure 2:**
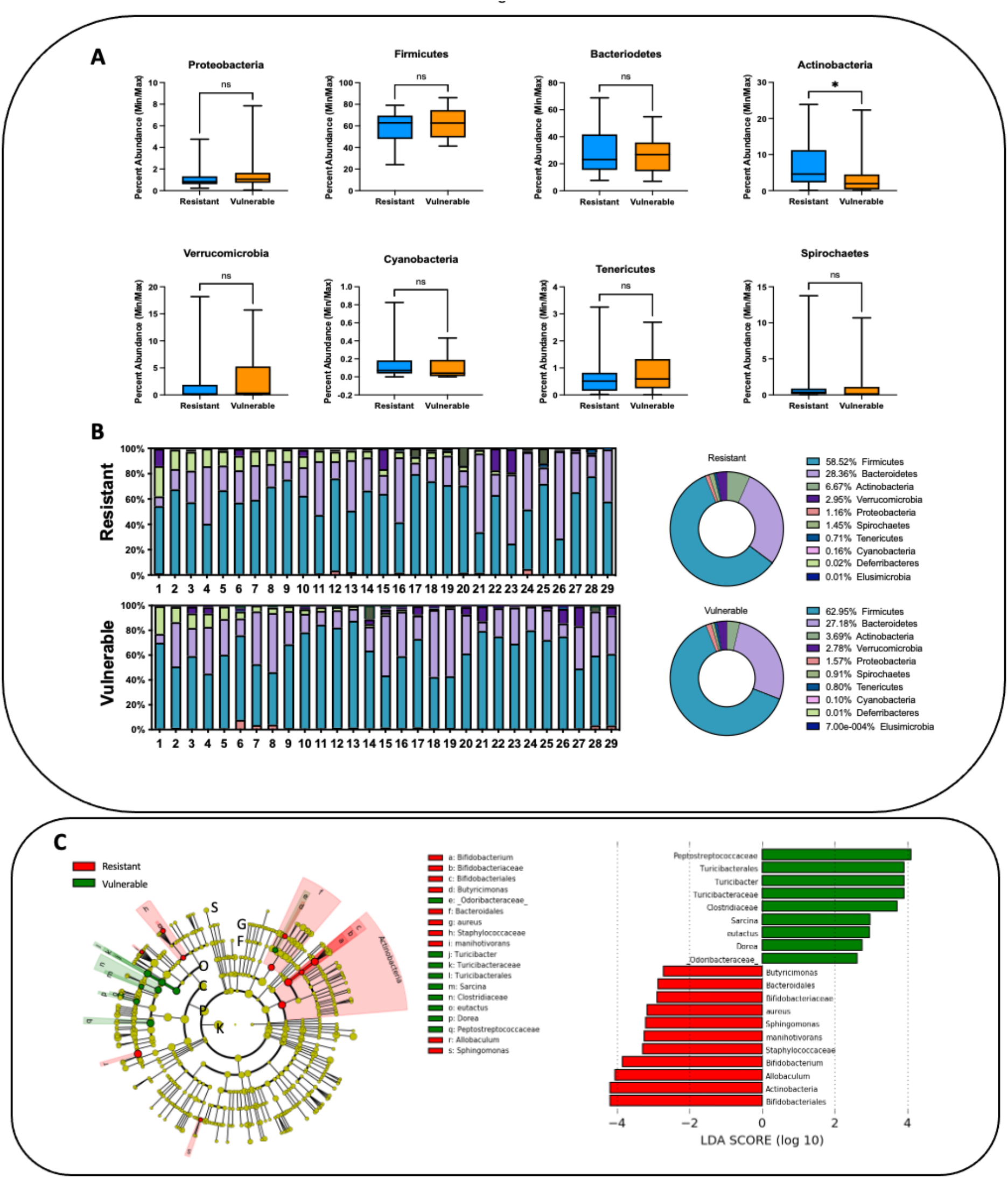
Phylum analysis of Resistant versus Vulnerable animals at baseline before drug exposure. **A)** Relative abundance of Proteobacteria (p=0.239), Firmicutes (p=0.307), Bacteroidetes (p>0.999), Verrucomicrobia (p=0.0778), Cyanobacteria (p=0.0228), Tenericutes (p=0.610), and Spiroachetes (p=0.245) were all not elevated in either group; Actinobacteria relative abundance is increased in the Resistant group compared to the Vulnerable group before drug exposure (p=0.025). B) Stacked bar plots demonstrating the relative abundance at the phylum level. C) Taxonomic cladograms obtained from linear discriminant analysis effect side (LEfSe) shows bacterial constituents that are more abundant in either the Resistant (red) or Vulnerable (green) groups. Linear discriminant effect size of the cladogram is plotted as bar charts to the right.

### Resistant and Vulnerable Populations Exhibit Differences in Microbiome Composition after Long-Access Cocaine Exposure

To assess the impact of long access cocaine self-administration sessions on the microbiome, we compared Resistant and Vulnerable populations at the phylum level. Again, Resistant and Vulnerable populations exhibited distinct microbial profiles in the Actinobacteria phylum (p = 0.030, Mann-Whitney U = 299) but also in the Tenericutes phylum (p = 0.032, Mann-Whitney U = 413.5). The Proteobacteria (p=0.5890), Firmicutes (p = 0.914), Bacteroidetes (p = 0.865), Verrucomicrobia (p=0.890), Cyanobacteria (p = 0.092), and Spirochaetes (p = 0.917) phyla were not differentially abundant between the two groups. Stacked bar charts shows the representative individuals from each group **(Figure 3B)**, and donut plots show the average relative abundance of each group. According to the LEfSe results plotted in the cladogram in **Figure 3C**, Vulnerable animals exhibited higher abundances at the phylum Defferibacteres and the class RF3 and Enterobacteriales, the order Defferibacterales, Clostridia, Klebsiella, and ML*615J_28*, the family, *Enterobacteriales, Ruminococcaceae*, the genus *Clostridium (unidentified) Ruminococcus, Roseburia, Blautia*, and *Facklamia*, and the species *Neonatale, Bromii, Oxytoca, Fragilis*, and *Schaedleri*. The LDA scores are plotted in the bar chart to the left.

**Figure 3:**
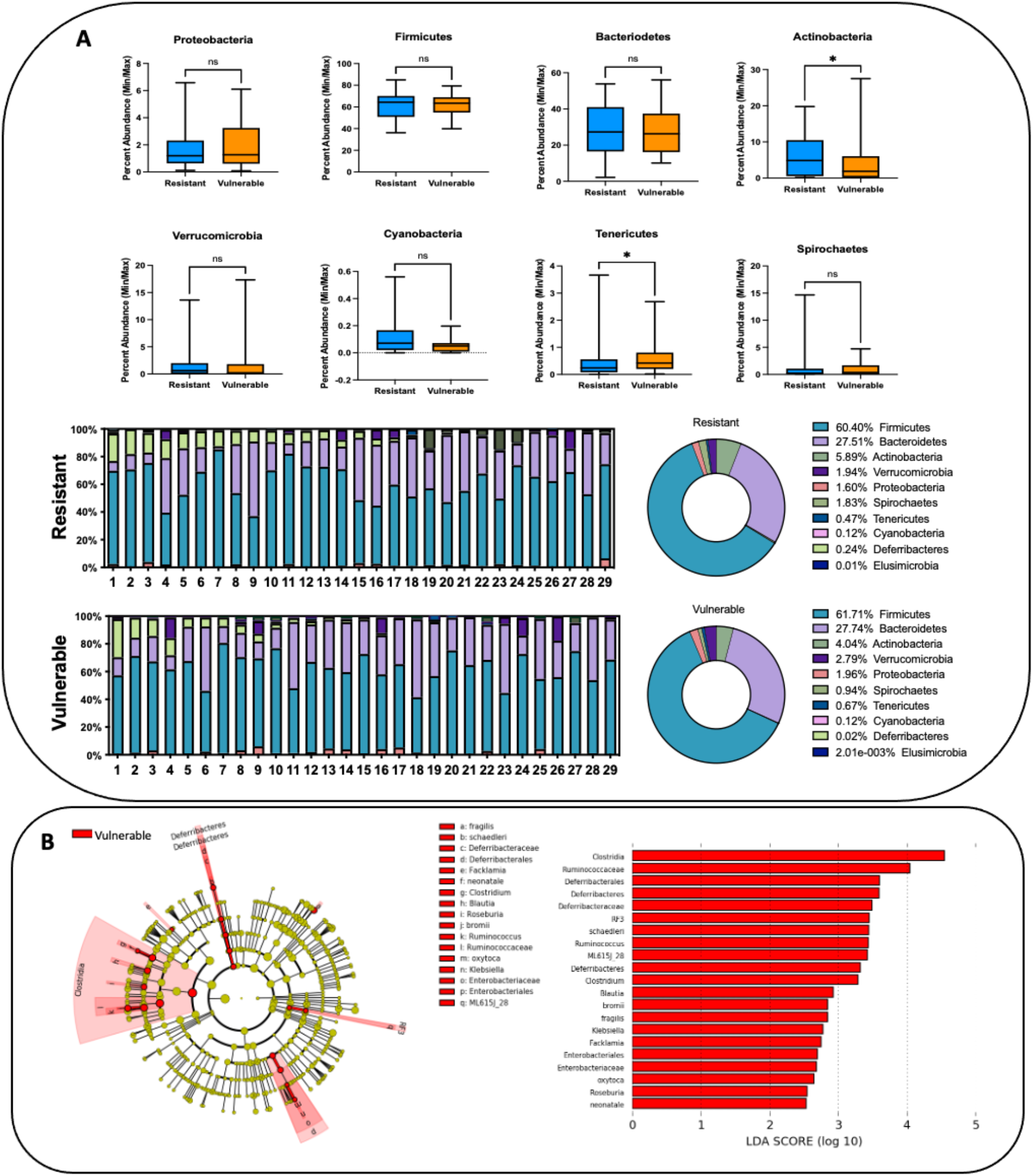
Phylum analysis of Resistant versus Vulnerable animals after long access cocaine exposure. **A)** Resistant and Vulnerable populations exhibited distinct microbial profiles in the Actinobacteria phylum (p = 0.030, Mann-Whitney U = 299) but also in the Tenericutes phylum (p = 0.032, Mann-Whitney U = 413.5). The Proteobacteria (p=0.5890), Firmicutes (p = 0.914), Bacteroidetes (p = 0.865), Verrucomicrobia (p=0.890), Cyanobacteria (p = 0.092), and Spirochaetes (p = 0.917) phyla were not differentially abundant between the two groups. **B)** Stacked bar plots demonstrate the relative abundance at the phylum level. **C)** Taxonomic cladograms obtained from linear discriminant analysis effect side (LEfSe) shows bacterial constituents that are more abundant in either the Resistant (red) or Vulnerable (green) groups. Linear discriminant effect size of the cladogram is plotted as bar charts to the right.

### PICRUSt analysis reveals alterations in SCFA processing in the Vulnerable group

Figure 4 highlights two KEGGs identified to be - K15833: Formate Dehyrogenlyase (p=0.045, Mann-Whitney U = 293.5) and K19709 (acetate CoA-transferase) (p=0.044, t=2.058) which are upregulated in the Vulnerable group compared to the Resistant group at the LgA timepoint. These two enzymes are involved in formate production and butyrate production.

### Sex differences reveal increased Actinobacteria at the phylum level in Male HS Rats

Sex as a biological variable is an important addition to our analysis. Similar to the stratification of groups with the addiction index, we split the male and female groups and measured rewards over time at the ShA and LgA timepoints **(Figure 5A)**. According to the two-way ANOVA there was no main effect difference between males and females at the ShA timepoint, but there was a significant effect of time (p < 0.001, F_(9,300)=_5.853), and an effect of group (p < 0.001, F_(1, 260)_ =18.2. According to the two-way ANOVA after the LgA timepoint there was a significant main effect (time x sex) (p=0.003, F_(13, 334)_ =2.74), there was also an effect of time (p<0.001, F_(13,390)_ =9.195) and an effect of sex (p=0.045, F_(1,30)=_4.365. The Bonferroni post-hoc revealed a significant difference between female and male rewards at day 12 (p=0.018) and day 14 (p=0.035). A student’s t-test of the progressive ratio at all three timepoints revealed no significant differences between males and females; however, there was a trend toward significance at PR2 (p=0.052) with females taking an increased number of rewards compared to males. Phylum analysis was compared at both Baseline **(Figure 5B)** and following Long Access **(Figure 5C)**. Males exhibited higher relative abundance of Actinobacteria (p<0.001, t=3.491) and lower relative abundance of Tenericutes (p=0.024, t=2.327) and Spirochetes (p=0.034, t=2.710) at the baseline timepoint. Males also exhibited higher relative abundance of Spirochetes (p=0.034, t=2.710) following Long Access, as well as a trend toward increased relative abundance of Cyanobacteria (p=0.061). The difference observed in Tenericutes at the baseline timepoint was not observed after long access (p=0.450). Additionally, Actinobacteria was no longer at higher abundance (p=0.469) in males compared to females at the Long Access timepoint. At baseline, the LEfSe cladogram demonstrates higher abundance of Actinobacteria in males at the phylum, order (Bifidobacteriales), and species (*Bifidobacterium)* levels compared to females. Males also exhibited higher abundance of deltaproteobacteria (order, *desulfrovibrionales*, and higher abundance of the species *Veillonellacea*. Females exhibited higher abundance at the order level (Ruminococcacea), and species level (*Marvinbryantia* and *Clostridium*) belonging to the phylum Firmicutes and increased abundance in the species and *Holdemania* which belongs to the phylum Firmicutes and order *Erysipelotrichales*. At the long access timepoint, males exhibited several of the same elevated features as baseline – *Marvinbryantia, Ruminococcaceae*, and *Clostridium*, with the addition of *SMB53*, which is a member of the family *Clostridiaceae*. Females exhibited increased abundance in the phylum *Spriochaetes* at all levels, with increased abundance in *Spriochaetacea, Spirochaetales*, and *Treponema*. The LDA scores are plotted to the right of each cladogram.

**Figure 4:**
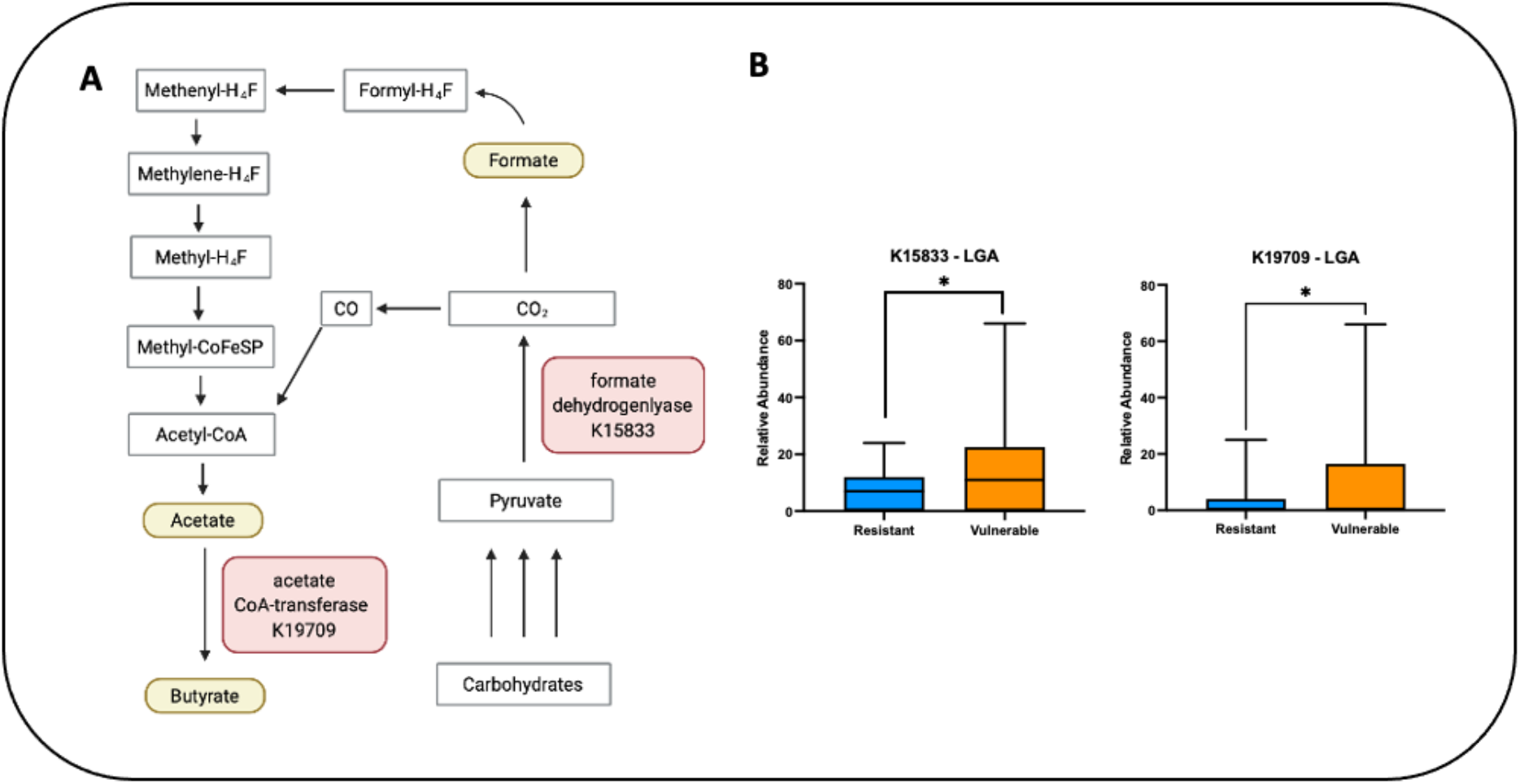
PICRUSt analysis after drug intake reveals increased abundance of KEGG hits related to enzymes involved in formate and butyrate production. **A)** Diagram of the location of the KEGG hits in the butyrate and formate production pathway. **B)** K15833 (Formate dehydrogenase) (p=0.045, Mann-Whitney U = 293.5) and K19709 (Acetate CoA-transferase) (p=0.044, t=2.058) gene content is more abundant in the Vulnerable group compared to the Resistant group.

**Figure 5:**
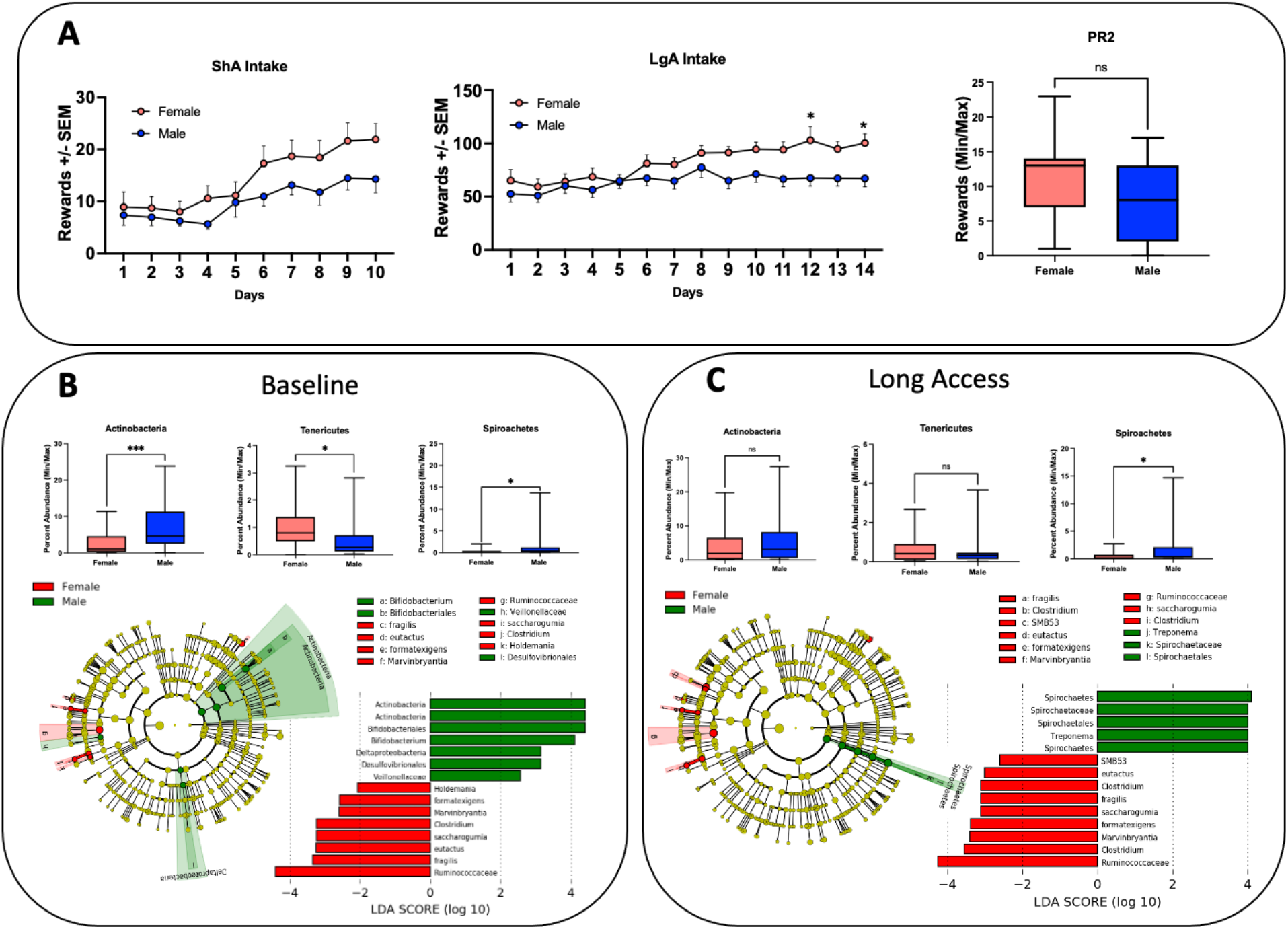
Phylum and behavioral analysis of males versus female animals after baseline and long access cocaine exposure. **A)** According to the two-way ANOVA there was no main effect difference between males and females at the ShA timepoint, but there was a significant effect of time (p < 0.001, F_(9,300)=_5.853), and an effect of group (p < 0.001, F_(1, 260)_ =18.2. According to the two-way ANOVA after the LgA timepoint there was a significant main effect (time x sex) (p=0.003, F_(13, 334)_ =2.74), there was also an effect of time (p<0.001, F_(13,390)_ =9.195) and an effect of sex (p=0.045, F_(1,30)=_4.365. The Bonferroni post-hoc revealed a significant difference between female and male rewards at day 12 (p=0.018) and day 14 (p=0.035). A student’s t-test of the progressive ratio at all three timepoints revealed no significant differences between males and females; however, there was a trend toward significance at PR2 (p=0.052) **B)** Baseline phylum analysis identified males exhibited higher relative abundance of Actinobacteria (p<0.001, t=3.491) and lower relative abundance of Tenericutes (p=0.024, t=2.327) and Spirochetes (p=0.034, t=2.710) at the baseline timepoint. **C)** Phylum analysis after long access showed that males also exhibited higher relative abundance of Spirochetes (p=0.034, t=2.710) following Long Access, as well as a trend toward increased relative abundance of Cyanobacteria (p=0.061). The difference observed in Tenericutes at the baseline timepoint was not observed after long access (p=0.450). Actinobacteria was no longer at higher abundance (p=0.469) in males. The taxonomic cladogram obtained from linear discriminant analysis effect side (LEfSe) shows bacterial constituents that are more abundant in either the female (red) or male (green) group. Linear discriminant effect size of the cladogram is plotted as bar charts to the right.

## Discussion

Cocaine use remains a public health problem with no medications, few available treatments (Kampman, 2019), and increasing overdose deaths. Recent developments in the microbiome space have highlighted the importance of microbes in drug taking behaviors (Kiraly et al., 2016a; Meckel and Kiraly, 2019; Simpson et al., 2020b). We tested the hypothesis that resistant individuals may possess protective microbiome constituents compared to those that are vulnerable to escalating intake of cocaine. A median split by addiction index allowed for the stratification of Resistant and Vulnerable individuals that reflected intake, compulsivity, and motivation. We also examined sex as a biological variable to determine if observed differences in intake and microbiome profile were related to sex. Sex emerged as a potential predictive indicator of drug intake and resilience. Males displayed increased ratios of protective microbes, while females expressed fewer protective microbes and consumed more cocaine as the study progressed. When subdivided by addiction index, Resistant individuals had higher abundance of beneficial microbiome constituents compared to the Vulnerable group. The behavioral stratification by addiction index in the HS strain of rat allows for better comparison of addiction liability in a strain with significant variation in drug taking behavior. Resistant animals exhibited decreased intake at ShA and LgA timepoints, decreased progressive ratio responding, which models motivation to obtain drug (Stafford et al., 1998), and decreased levels of intake in a model of compulsivity (shock) (Johnson and Kenny, 2010). This approach best models the multi-dimensional behavior observed in humans.

Comparing the microbiome of Vulnerable and Resistant animals at baseline highlighted some differences in microbiota composition. For instance, Actinobacteria, the third largest phyla were increased in Resistant individuals. Upon further inspection at deeper levels, *Bifidobacterium* was increased in the Resistant group relative to the Vulnerable group. *Bifidobacterium* has been shown to be enriched in individuals that were protected from liver cirrhosis in alcohol dependent individuals (Dubinkina et al., 2017), and supplementation of *Bifidobacterium* was associated with improvement in alcohol-induced liver injury (Kirpich et al., 2008). *Bifidobacterium* has also confers resilience in a model of chronic social defeat stress, and administration of *Bifidobacterium* has been hypothesized to minimize relapse of depression induced by inflammation/stress (Yang et al., 2017). *Butyricimonas*, a species of butyrate producing bacteria was also increased in the resistant group relative to the vulnerable group (Barcenilla et al., 2000). *Butyricimonas* express genes that code for acetyl-CoA synthetase [EC:6.2.1.1] which irreversibly converts acetate to acetyl-CoA (Leclercq et al., 2020). Disruption of SCFA production has been demonstrated to lower the threshold of reward in cocaine (Kiraly et al., 2016a). Vulnerable animals exhibited increased *Turcibacter, Clostridia, Peptostreptococcaceae* at the order and genus level. Similarly, mice exposed to cocaine have been demonstrated to have increased levels of *Turcibacter* (Chivero et al., 2019). Constituents of *Turcibacter* and *Clostridia* have been demonstrated to interact with the serotonergic system in the gut, and to increase upon elevating levels of serotonin either by oral supplementation of genetic deficiency of the host serotonin transporter (Fung et al., 2019). *Peptostreptococcaceae* was more abundantly expressed in animals that had high conditioned place preference for morphine (Zhang et al., 2020), and increases in *Peptostreptococcaceae* have been observed in chronic schizophrenia patients (Liu et al., 2021).

In contrast to the microbiome compositions observed at baseline, after both short and long access cocaine exposure, Resistant animals had several beneficial abundant constituents of the class *Clostridia* and *RF3*, the family *Ruminococcus*, and the genus *Roseburia, Clostridium*, and *Enterobacteriaceae*. The genus *Roseburia* is a well-known producer of Butyrate and is associated with positive health outcomes (Tamanai-Shacoori et al., 2017). In other drugs of abuse, the depletion of *Roseburia* is associated with increase alcohol intake (Seo et al., 2020), and decreased *Roseburia* may relate to increased gut permeability/inflammation and dysregulation of bile acids (Gicquelais et al., 2020). A higher abundance of butyrate producing *Clostridia* in the Resistant animals is also protective as depletion of *Clostridia* can increase aerobic expansion of opportunist microbes and increase inflammation (Rivera-Chavez et al., 2016).

Among the most crucial components of this study is the potential to identify microbial constituents of cocaine resilience. A broad theme in differentially abundant microbes is the production and processing of SCFA in resistant individuals before cocaine exposure. PICRUSt analysis identified two KEGGs following LgA that demonstrated upregulation of pathways related to SCFA production. K15833: Formate Dehyrogenlyase and K19709 (acetate CoA-transferase) are upregulated in the Vulnerable group compared to the Resistant group at the LgA timepoint. It is likely that with a reduction in butyrate producing bacteria compared to the Resistant group that the Vulnerable microbiota are compensating for a decrease in available SCFA. Dysregulation of available SCFAs has been demonstrated to play a role in the alterations of cocaine CPP and lower the reward threshold for cocaine (Kiraly et al., 2016a).

In addition to addiction index, observed differences in intake based on sex is vital in the development of effective novel treatments. Even though men are more likely to use illicit drugs than women, women are just as likely to develop substance use disorders. It has been hypothesized that women may also have increased susceptibility to relapse and craving compared their male counterparts (Robbins et al., 1999; Hitschfeld et al., 2015). In this study, cocaine intake was higher in females at LgA than males toward the end of the testing phase. While progressive ratio was not significantly increased in females compared to males (PR2), it trended toward significance (p=0.052). Baseline microbiome differences were observed at the phylum level, including increased abundance of Actinobacteria in males compared to females, and decreased Tenericutes and Spiroachetes in males compared to females. Differences in Actinobacteria and Tenericutes were lost after long access exposure to cocaine, and a reversal in abundance in Spiroachetes was observed in males at the LgA timepoint. Beneficial butyrate producers were more abundant in males compared to females prior to drug exposure, which was reversed after drug exposure. After drug exposure, males exhibited increased abundance in known opportunists (Spiroachetes, Treponema) compared to females. While not causal in nature, potential differential microbiome constituents may play a role in the observed behavioral differences in this study. Further manipulations are needed to tease apart the contributions of sex to the microbiome in the context of drug taking behavior.

As mentioned above, there are a few potential shortcomings to consider from our study. Profiling fecal samples constrains the observed microbes to only the final snapshot of microbes present in the colon; however, repeated sampling throughout the gut without significantly perturbing the system is still difficult to accomplish and the ability to sample longitudinally outweighs the drawback (Stanley et al., 2015). Another consideration is the observational nature of our study we cannot attribute causality to the makeup of the gut microbiome. Further functional studies to follow up on our findings are needed.

Despite the limitations of the present study, the gut-brain axis has emerged as a contributor to substance use disorders, including cocaine vulnerability/resilience. The present study explores this hypothesis further by identifying microbial populations that are associated with resilience to cocaine escalation by a multi-dimensional addiction index as well as contributions of sex to differences in the microbiome profile. Furthermore, analysis of predicted gene function using PICRUSt identified potential compensation in SCFA production in the Vulnerable animal group, aligning with previous studies that demonstrated depletion of SCFA producing bacteria lowered the reward threshold for cocaine in models of CPP. Furthermore, in contrast to the Vulnerable group, the Resistant group exhibited increased abundances of known butyrate producing microbes prior to drug exposure. Efforts to identify microbes and downstream metabolic pathways that may confer resistance will enable the development of potential treatments for CUD that are applicable to both sexes.

## Acknowledgements

We thank the Cocaine Biobank for access to samples and the Preclinical Addiction Research Consortium for support.

